# Genome-resolved insights into microbial diversity and elemental cycling in Winogradsky columns

**DOI:** 10.64898/2026.08.29.748020

**Authors:** Stefan P. Anthopoulos, Kaitlyn P. Boutwell, Griffin T. Deans, Michael J. Glinski, Zihan Zhong, Khulan Byambasuren, Alexis J. Miskelly, Puja Shrestha, Beryl Braden, Rita Faivre-Nigro, Kaelin Feliu, Emily Garlock, Asha G. Hotaling, Melia G. Kanaovicz, Bryce E. Manning, Keith McGill, Samantha Phoenix, Daniel Ryu, Jackson L. Solfrian, Carlos A. Rodriguez-Bornot, Jean Yang, Jennifer L. Goff

## Abstract

Winogradsky columns are a classic model ecosystem for studying microbial biogeochemistry across steep gradients of oxygen and sulfide. They also remain widely used in microbiology education, introducing generations of students to microbial diversity. Yet, the genomic potential of their microbial communities remains uncharacterized. Here, we applied shotgun metagenomic sequencing to a Winogradsky column community at multiple depths, yielding 20 metagenome-assembled genomes (MAGs) representing diverse, largely uncultivated taxa. Genome-resolved analyses revealed metabolically diverse oxygenic and anoxygenic phototrophs that could potentially contribute to carbon and nitrogen fixation across all layers of the column. Most of these phototrophs also encoded one or more pathways for sulfur oxidation, which we speculated may support both energy conservation and/or sulfide detoxification by these populations. Complex carbon degradation capacity was also widespread across the MAGs, suggestive of the potential for the transformation of the column’s amended organic matter (shredded coffee filters) into smaller depolymerization products and, through fermentation, organic acids. Together, these findings reveal how distinct microbial guilds might partition interconnected carbon, sulfur, and nitrogen transformations within redox-stratified systems.

**IMPACT STATEMENT:** Although Winogradsky columns have served as both an enrichment platform and teaching staple for over a century, the microorganisms associated with their characteristic redox stratification have not been characterized at the genomic level. Here, we reconstruct genomes from a column community, showing how largely uncultivated taxa divide carbon, sulfur, and nitrogen cycling across the column’s gradients of oxygen and sulfide. This work transforms a widely used—yet genomically opaque—system into a now genome-resolved model for studying microbial community metabolism, with value for both researchers investigating redox-stratified environments and educators introducing students to microbial diversity and metabolism.

**DATA SUMMARY:** Metagenome-assembled genomes and read libraries are available in the following KBase narrative: https://doi.org/10.25982/266974.4/3412610. Both individual and merged (*i.e.,* the library used for the co-assembly) read libraries are deposited in the Sequence Read Archive: [SRR40126710-14](https://www.ncbi.nlm.nih.gov/Traces/study/?acc=SRP726339&o=acc_s%3Aa) and [SRR40144435](https://www.ncbi.nlm.nih.gov/Traces/study/?acc=SAMN62374358). MAGs with greater than 90% completion were also deposited with NCBI with the BioSample numbers: **SAMN62641021-31 [**REVIEW NOTE: NCBI IS STILL PROCESSING THESE SUBMISSIONS BUT THE MAGS ARE AVAILABLE THROUGH THE KBASE **NARRATIVE DOI**]. Note that we use GTDB taxonomy throughout the paper, which does not always align with NCBI taxonomy, though the strain names are consistent. All sequencing data deposited with NCBI are associated with the BioProject: PRJNA1511471

## INTRODUCTION

Sergei Winogradsky is considered one of the founders of modern microbial ecology (1), uncovering the role of microorganisms in the biogeochemical cycles of sulfur and nitrogen (1). One of Winogradsky’s enduring contributions was the development of an enrichment-based cultivation approach for isolating environmental microorganisms that were otherwise recalcitrant to contemporaneous laboratory cultivation methods (2). In 1888, he described isolating *Beggiotoa* by incubating mud and plant material in a deep jar amended with gypsum—observing visible stratification as distinct microbial populations developed along opposing gradients of oxygen, decreasing from top to bottom, and sulfide, increasing from bottom to top (3–5). Subsequent researchers expanded upon this strategy, which became known as a “Winogradsky column” (6–9).

Today, Winogradsky columns remain useful for enriching metabolically diverse microorganisms, including arsenic-oxidizing (10), acetaminophen-degrading (11), and plastic-degrading bacteria (12–16), as well as for studying rare microbial eukaryotes (17) and microbial community responses to varying environmental conditions (5, 18–22). Perhaps their most enduring application may be as a teaching tool for introducing concepts in microbial diversity, ecology, and metabolism (23–25).

Despite longstanding usage, relatively little is known about the full metabolic potential of Winogradsky column communities. Existing studies have relied exclusively on amplicon sequencing (4, 19, 26–29), which provides limited insight into the functional potential of microbial communities. Yet the features that make these columns compelling from an educational standpoint (*i.e.,* steep redox gradients that reproducibly organize a stratified microbial community) also make them a relevant model for certain natural environments, including sediments (30, 31), microbial mats (32, 33), and redox-stratified water columns (34–36). Thus, applying genome-resolved metagenomics to Winogradsky columns can enhance our understanding of uncultivated microbial diversity, metabolic potential, and biogeochemistry in redox-stratified systems more generally.

Here we report, to our knowledge, the first metagenomic analysis of a Winogradsky column using an enrichment strategy conceptually similar to Winogradsky’s original *Beggiatoa* enrichment and widely described in the teaching literature (23, 37–39). Reflecting the column’s educational legacy, the work was conducted across two courses at SUNY College of Environmental Science and Forestry. Columns were constructed by graduate landscape architecture students to visualize the microbial dimensions of landscape ecology, and DNA from two columns was subsequently analyzed by undergraduate and graduate students in a Microbial Biochemistry course.

## MATERIALS AND METHODS

### Winogradsky column establishment and incubation

Water collected at a decorative pond at the Stone Quarry Hill Art Park (Cazenovia, NY, 42.911401265775744, -75.83483059444166) was used as the column inoculum. Approximately 2-3 grams of shredded coffee filters were added to the bottom of each column (a 1.4 L glass cylinder), along with ∼2 grams of Plaster of Paris (calcium sulfate hemihydrate) as a sulfur source and ∼2 grams of crushed chalk as a carbonate buffer (calcium carbonate). Sand (∼400 grams) was added to each column, followed by pond water to fill each column about 2/3 of the total volume. The columns were incubated in a Percival Scientific incubator with constant fluorescent lighting at room temperature for three months, sealed with a gas-permeable membrane.

### DNA extraction and sequencing

Triplicate sediment samples were collected from three heights (sediment surface biofilm, shallow subsurface sediments, and deep subsurface sediments) along the edges of the column **(Fig. S1),** representing the column’s distinct visible layers (**Supporting Information)**. DNA extraction and Illumina short-read sequencing methods are described in the **Supporting Information.** The samples that yielded sufficient DNA for subsequent sequencing were as follows: sediment surface biofilms in two different columns (one sample from column 1 and two samples from column 2), a shallow subsurface sample (∼2.5 cm below the surface) from column 2, and a deep subsurface sample (∼7.5 cm below the surface) from column 2.

### Metagenomic assembly, binning, and annotation

Details of our computational pipelines are in the **Supporting Information**. Briefly, the five metagenomic libraries were assembled and binned both separately and as a single co-assembly to maximize MAG recovery. We used the MIMAG standards (39) to report MAG quality, retaining all medium- and high-quality draft genomes for further analysis, along with a few borderline cases, as detailed in the **Supporting Information**. Our downstream analyses are detailed in the **Supporting Information**.

## RESULTS AND DISCUSSION

### Winogradsky columns support novel bacterial lineages

Metagenomic sequencing of the Winogradsky column samples (**Fig. S1**) yielded 14,648,035,672 bp of DNA, which was assembled and binned into metagenome-assembled genomes (MAGs). We retained 18 MAGs with ≥50% completion and <10% contamination (**Table 1**). Of these, 10 had >90% completion and <5% contamination; however, only one (*Paceibacteria* Bin.003_MG) can be described as high-quality, as the others lacked a full suite of 5S, 16S, and 23S rRNA genes. We also retained two borderline medium-quality MAGs: one MAG (*Chromatiaceae* Bin.008_AM) with >40% completion and <5% contamination, as well as one (*Rhodopseudomonas* Bin.015_MK) with >90% completion and <20% contamination (**Table 1**). Significant chimerism (**Table S1**) was not detected in this latter MAG (41), suggesting that it may represent two closely related strains collapsing into a single bin.

**Table 1.** Bin Statistics. Detailed bin statistics of MAGs retained for this analysis. %Complete. = %Completeness; %Contam = % Contamination. High-quality draft genomes were defined as: >90% completion, <5% contamination, and the presence of the 23S, 16S, and 5S rRNA genes and at least 18 tRNA genes. Medium-quality draft genomes were defined as: ≥50% completion and <10% contamination.

| Bin Name | Contigs | Length (bp) | %GC | %Complete. | %Contam. | unique tRNAs | 5S rRNAs | 16S rRNAs | 23S rRNAs | MIMAG Quality |
| --- | --- | --- | --- | --- | --- | --- | --- | --- | --- | --- |
| Bryobacteraceae_Bin.002_AH | 681 | 3,333,200 | 62.95 | 76.13 | 0.00 | 31 | 0 | 0 | 0 | Medium |
| Bacteroidia_Bin.008_KM | 908 | 4,088,358 | 57.06 | 52.59 | 0.00 | 21 | 0 | 1 | 0 | Medium |
| Chlorobaculum_Bin.017_DR | 23 | 2,581,485 | 57.60 | 98.90 | 1.10 | 45 | 0 | 0 | 0 | Medium |
| Chloroherpetonaceae_Bin.002_PS | 499 | 2,562,123 | 48.78 | 78.02 | 2.73 | 39 | 1 | 1 | 0 | Medium |
| Chloroflexaceae_Bin.020_RN | 144 | 3,541,111 | 48.97 | 91.98 | 0.00 | 42 | 0 | 2 | 0 | Medium |
| Cyanobacteriales_Bin.017_ZZ | 56 | 5,791,177 | 54.60 | 98.89 | 1.00 | 71 | 0 | 0 | 0 | Medium |
| Hassalia_Bin.018_JS | 1036 | 5,390,677 | 41.12 | 75.89 | 0.13 | 29 | 1 | 0 | 0 | Medium |
| Kamptonema_Bin.017_EG | 501 | 6,587,525 | 42.06 | 91.14 | 0.33 | 69 | 0 | 0 | 0 | Medium |
| Gemmatimonadales_Bin.002_JG | 1010 | 2,765,040 | 66.67 | 57.66 | 9.77 | 34 | 1 | 2 | 0 | Medium |
| Moranbacterales_Bin.008_KB | 29 | 1,151,931 | 54.63 | 93.02 | 4.65 | 48 | 1 | 1 | 0 | Medium |
| Paceibacteria_Bin.003_MG | 15 | 947,620 | 57.87 | 90.70 | 0.00 | 44 | 1 | 1 | 1 | High |
| Patescibacteria_Bin.011_SA | 146 | 1,167,625 | 34.60 | 67.44 | 9.30 | 38 | 1 | 0 | 0 | Medium |
| Phycisphaerales_Bin.012_GD | 130 | 3,115,706 | 65.80 | 96.02 | 0.10 | 43 | 0 | 2 | 1 | Medium |
| Chromatiaceae_Bin.008_AM | 462 | 1,789,649 | 66.01 | 41.45 | 0.29 | 27 | 0 | 0 | 0 | Low |
| Phaeospirillum_Bin.004_BAB | 308 | 3,798,130 | 69.22 | 93.61 | 1.74 | 39 | 0 | 0 | 0 | Medium |
| Phaeospirillum_Bin.015_KB | 100 | 3,321,753 | 66.30 | 98.26 | 4.98 | 42 | 0 | 0 | 0 | Medium |
| Phaeospirillum_Bin.019_SP | 123 | 3,282,643 | 66.02 | 95.71 | 2.74 | 44 | 0 | 0 | 0 | Medium |
| Rhodopseudomonas_Bin.015_MK | 333 | 4,807,028 | 64.25 | 97.82 | 17.05 | 56 | 0 | 1 | 0 | Low |
| Spongiibacteraceae_Bin.006_AM | 431 | 1,871,631 | 47.4 | 56.43 | 0.50 | 17 | 0 | 0 | 0 | Medium |
| Steroidobacteraceae_Bin.004_BM | 643 | 2,381,344 | 66.82 | 57.24 | 7.41 | 14 | 0 | 0 | 0 | Medium |

All MAGs were bacterial, spanning eight phyla (**Table 2**). Six of the MAGs (*Bryobacteraceae* Bin.002 AH, *Chloroherpetonaceae* Bin.002_PS, *Patescibacteria* Bin.011_SA, *Phycisphaerales* Bin.012_GD, *Chromatiaceae* Bin.008_AM, and *Spongiibacteraceae* Bin.006_AM) represented novel genera. Additionally, three were from undescribed families (*Cyanobacteriales* Bin.017_ZZ, *Gemmatimonadales* Bin.002_JG, *Phycisphaerales* Bin.012_GD), two were from undescribed orders (*Paceibacteria* Bin.003_MG, *Bacteroidia* Bin.008_KM), and one of the MAGs (*Patescibacteria* Bin.011_SA) was from an undescribed class. Interestingly, we recovered three *Phaeospirillum* MAGs (Bin.004_BAB, Bin.015_KB, Bin.019_SP). As these MAGs had ANIs of 81-83% among themselves, all three were retained as separate species for further analysis. These 20 MAGs accounted for 11% to 70% of the total microbial communities in each sample (**Fig. S2**, **Table S2**). We next examined the biogeographic distribution of the MAGs along the Winogradsky column’s opposing gradients of oxygen (decreasing with depth) and sulfide (increasing with depth) (24, 41) to determine how their spatial distributions corresponded to their metabolic potential.

**Table 2.** Bin Taxonomy. GTDB classification for all MAGs retained for analysis. All MAGs are members of the domain Bacteria. NA = no designation could be made to that taxonomic level.

| Bin Name | Phylum | Class | Order | Family | Genus |
| --- | --- | --- | --- | --- | --- |
| Bryobacteraceae_Bin.002_AH | Acidobacteriota | Terriglobia | Bryobacterales | Bryobacteraceae | NA |
| Bacteroidia_Bin.008_KM | Bacteroidota | Bacteroidia | J057 | JADKCL01 | JADKCL01 |
| Chlorobaculum_Bin.017_DR | Bacteroidota | Chlorobia | Chlorobiales | Chlorobiaceae | Chlorobaculum |
| Chloroherpetonaceae_Bin.002_PS | Bacteroidota | Chlorobia | Chlorobiales | Chloroherpetonaceae | NA |
| Chloroflexaceae_Bin.020_RN | Chloroflexota | Chloroflexia | rexales | Chloroflexaceae | UBA1466 |
| Cyanobacteriales_Bin.017_ZZ | Cyanobacteriota | Cyanobacteriia | Cyanobacteriales | FACHB-T130 | FACHB-831 |
| Hassalia_Bin.018_JS | Cyanobacteriota | Cyanobacteriia | Cyanobacteriales | Nostocaceae | Hassalia |
| Kamptonema_Bin.017_EG | Cyanobacteriota | Cyanobacteriia | Cyanobacteriales | Microcoleaceae | Kamptonema |
| Gemmatimonadales_Bin.002_JG | Gemmatimonadota | Gemmatimonadetes | Gemmatimonadales | GWC2-71-9 | JAGGAG01 |
| Moranbacterales_Bin.008_KB | Patescibacteria | Paceibacteria | Moranbacterales | UBA1568 | JAAXTX01 |
| Paceibacteria_Bin.003_MG | Patescibacteria | Paceibacteria | UBA9983_A | UBA2100 | UBA10103 |
| Patescibacteria_Bin.011_SA | Patescibacteria | JACMRA01 | JACMRA01 | JACMRA01 | NA |
| Phycisphaerales_Bin.012_GD | Planctomycetota | Phycisphaerae | Phycisphaerales | SM1A02 | NA |
| Chromatiaceae_Bin.008_AM | Pseudomonadota | Gammaproteobacteria | Chromatiales | Chromatiaceae | NA |
| Phaeospirillum_Bin.004_BAB | Pseudomonadota | Alphaproteobacteria | Rhodospirillales | Magnetospirillaceae | Phaeospirillum |
| Phaeospirillum_Bin.015_KB | Pseudomonadota | Alphaproteobacteria | Rhodospirillales | Magnetospirillaceae | Phaeospirillum |
| Phaeospirillum_Bin.019_SP | Pseudomonadota | Alphaproteobacteria | Rhodospirillales | Magnetospirillaceae | Phaeospirillum |
| Rhodopseudomonas_Bin.015_MK | Pseudomonadota | Alphaproteobacteria | Rhizobiales | Xanthobacteraceae | Rhodopseudomonas |
| Spongiibacteraceae_Bin.006_AM | Pseudomonadota | Gammaproteobacteria | Pseudomonadles | Spongiibacteraceae | NA |
| Steroidobacteraceae_Bin.004_BM | Pseudomonadota | Gammaproteobacteria | Steroidobacterales | Steroidobacteraceae | RPQJ01 |

### Diverse phototrophs perform carbon and nitrogen fixation at the top of the column

Phototrophic MAGs recovered from the Winogradsky columns fall into three groups: oxygenic phototrophs, Type I anoxygenic phototrophs, and Type II anoxygenic phototrophs. These phototrophs are predicted to provide both organic carbon and fixed nitrogen to the column community.

*Cyanobacteriales* Bin.017_ZZ, *Hassalia* Bin.018_JS, and *Kamptonema* Bin.017_EG are oxygenic phototrophs that likely contribute to oxygen accumulation in the upper column, as suggested by bubbles within the biofilms (**Fig. S1**), but are also predicted to respire aerobically in the dark (**Tables S3-S4**). They encode nearly complete PSI and PSII systems as well as the genes for carbon fixation via the Calvin cycle (*prk, rbcS, rbcL*) (**Tables S4-S5**). These MAGs were most abundant in the green-pigmented sediment surface biofilms (**Fig 1, Fig. S1**), where *Cyanobacteriales* Bin.017_ZZ dominated (29-37% relative abundance) (**Fig. S2**). This MAG remained the dominant member of the shallow subsurface sediments (29%), where we also observed green pigmentation (**Figs. S1-S2**), but declined in the deep subsurface sediments (0.2%) (**Fig. 1**). The carbon predicted to be fixed by these organisms can be transferred downward via cell lysis and extracellular release, serving as a carbon source for other microbes in deeper layers (43, 44).

**Figure 1.**
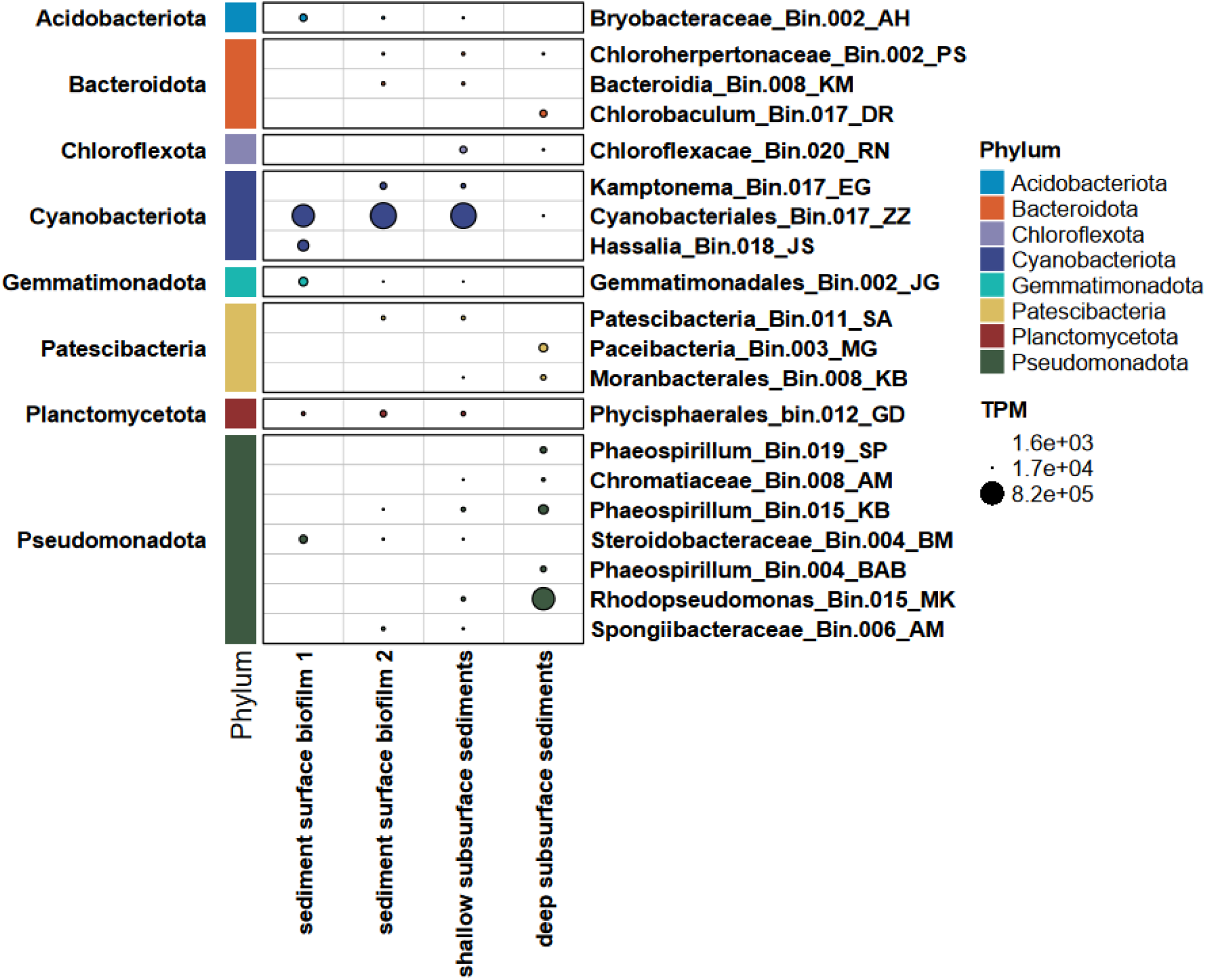
Abundance of each MAG across depths in the Winogradsky column. Read counts mapped to each MAG retained for analysis (described in the Materials & Methods) are normalized by “transcripts per kilobase million” (TPM). Images of the columns are shown in **Fig. S1**.

Type I anoxygenic phototrophs included the green sulfur bacteria (GSB) *Chlorobaculum* Bin.017_DR and *Chloroherpetonaceae* Bin.002_PS, encoding *pscA–D* (**Table S3**) and predicted to fix carbon through the reverse TCA cycle (*aclA, aclB*) (**Table S5**). Both occurred primarily in the sediments, with *Chlorobaculum* Bin.017_DR restricted to the deeper sediments (∼4% relative abundance, **Fig. 1**), where we observed a green pigmentation (**Fig. S1**)—consistent with the characteristic low-light, anoxic, sulfide-rich niche of GSB (45).

Diverse Type II anoxygenic phototrophs (encoding *pufL/pufM*) were also recovered (**Table S3**). Within the sediments, low to moderate abundances (∼0.1-7%) were observed of the green non-sulfur bacterium (GNSB) *Chloroflexaceae* Bin.020_RN; the purple sulfur bacterium (PSB) *Chromatiaceae* Bin.008_AM; and the purple non-sulfur bacteria (PNSB) *Phaeospirillum* MAGs Bin.004_BAB, Bin.015_KB, and Bin.019_SP (**Fig. 1, Fig. S1**). Of these, the latter four are predicted to fix carbon by the Calvin Cycle (**Table S5**).

The *Phaeospirillum* MAGs had distinct distributions within the column: Bin.004_BAB and Bin.019_SP were exclusively found at the bottom of the column, while Bin.015_KB was also identified in both the surface-associated biofilms and both sediment locations (**Fig. 1**). Notably, Bin.015_KB encodes a complete type III-B CRISPR-Cas system (absent from the other two strains) along with a type I-C system that was also present in Bin.019_SP, whereas no CRISPR-Cas system was detected in Bin.004_BAB. As phages have been previously suggested to structure Winogradsky column community dynamics (25), and because type III systems can counteract viral escape from type I systems (44, 45), we speculate that this broader CRISPR-Cas repertoire might have been one factor facilitating Bin.015_KB’s wider distribution in the column.

The PNSB *Rhodopseudomonas* Bin.015_MK was particularly abundant (∼43%) in the deep subsurface sediments, where we also observed a reddish-purple pigmentation in the column (**Fig. 1, Fig. S1-2**). Members of *Rhodopseudomonas* are metabolically versatile, capable of photoautotrophic, photoheterotrophic, chemoautotrophic, and chemoheterotrophic growth (52, 53). Indeed, *Rhodopseudomonas* Bin.015_MK appears capable of growth via all four metabolic modes (**Tables S3-S5**). We speculate that this metabolic flexibility may have facilitated its relative success in the column, enabling it to exploit both photosynthetic carbon fixation and heterotrophic reuse of organic carbon (48).

Finally, no bioavailable nitrogen was added to the column beyond that naturally present in the source water. Consistent with this apparent nitrogen limitation, nitrogen-fixation capacity is widespread (**Table S6**): MoFe-nitrogenase genes (*nif*) were detected in all phototrophic MAGs. Field studies of microbial mats have similarly detected *nifH* expression across *Cyanobacteriota* and purple and green anoxygenic phototrophs (49).

Alternative FeFe and VFe nitrogenases were also observed. *Chlorobaculum* Bin.017_DR encodes FeFe nitrogenase (*anf*). FeFe nitrogenase genes were also observed in all three *Phaeospirillum* MAGs, while the VFe nitrogenase (*vnf*) was observed in Bin.004_BAB. The VFe nitrogenase could provide Bin.004_BAB with an alternative mechanism for nitrogen fixation under molybdenum-limited conditions (50, 51), which is likely relevant to the bottom of the column where sulfide is expected to accumulate due to the sulfate amendments (52).

### Winogradsky columns enrich for diverse sulfur-oxidizing pathways

Diverse sulfur oxidation pathways were identified in MAGs distributed throughout the column (**Figs. 1-2**, **Table S7**), consistent with upward diffusion of sulfide from deeper, more reducing layers (59). These genes were exclusive to phototrophic MAGs, linking sulfur cycling to primary productivity. We propose that sulfur oxidation within the Winogradsky column serves dual purposes: (*1*) detoxification of excess sulfide and (*2*) electron donation to the phototrophic electron transport chain.

**Figure 2.**
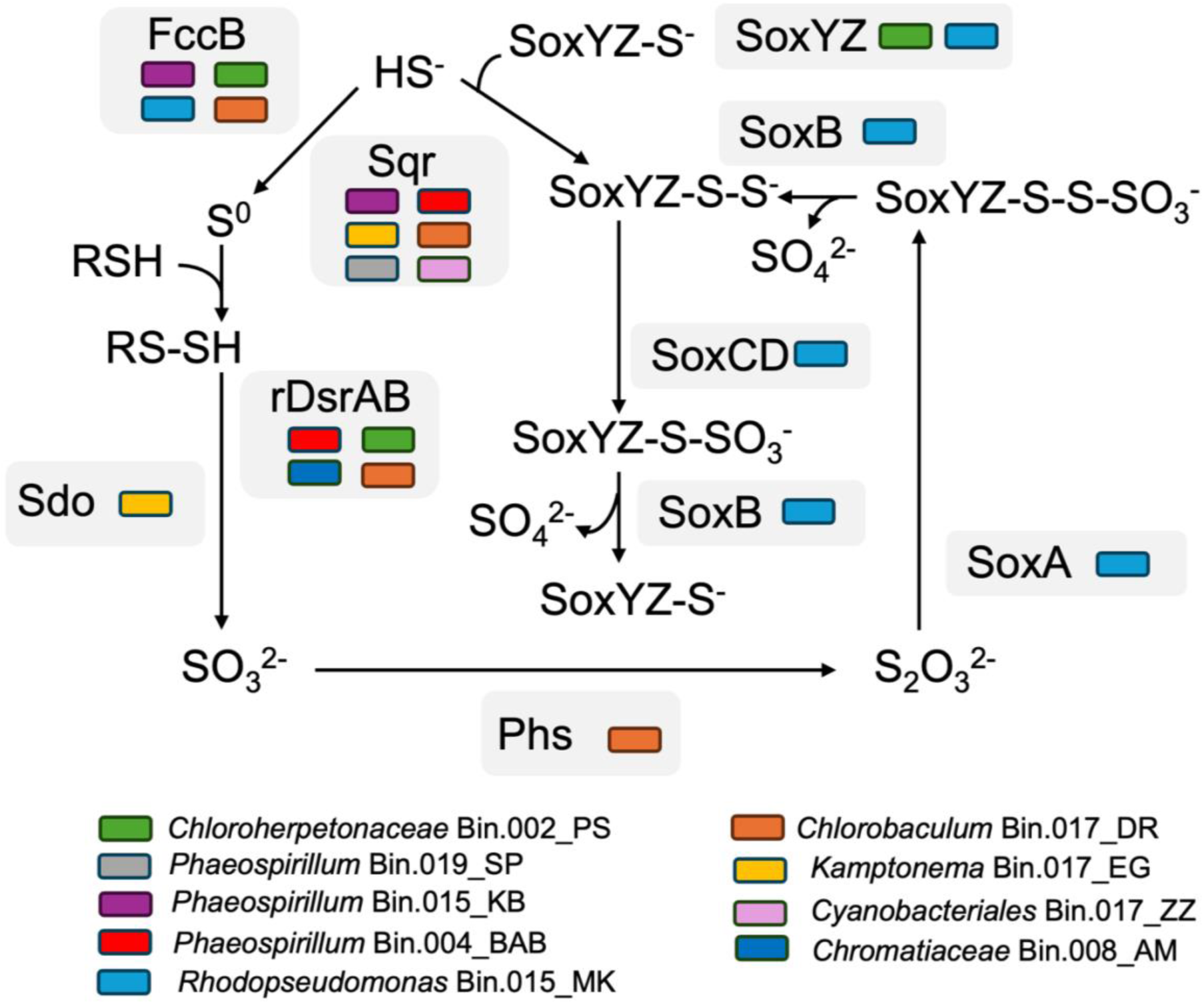
Sulfur oxidation pathways predicted in the Winogradsky column. The colored square next to each enzyme name indicates that the enzyme is predicted to be encoded by that MAG. Molecules shown: HS^-^ = sulfide, S^0^ = zero-valent sulfur, RSH = thiol, RS-SH = hydropersulfide, SO_3_^2-^ = sulfite, S_2_O_3_^2-^ = thiosulfate, SO_4_^2-^ = sulfate. Enzymes shown: FccB = sulfide dehydrogenase, Sqr = sulfide-quinone reductase, rDsrAB =reverse dissimilatory sulfite reductase, Phs = thiosulfate reductase,

Several of the phototrophic MAGs encode genes for the sulfide-oxidizing enzymes FccB (sulfide dehydrogenase) and Sqr (sulfide-quinone reductase), which produce zero-valent sulfur (ZVS). These include the PSB and GSB MAGs, the PNSB *Phaeospirillum* MAGs, and two of the oxygenic phototroph MAGs (*Cyanobacteriales* Bin.017_ZZ and *Kamptonema* Bin.017_EG (**Fig. 2**). Unlike PSB, phototrophic growth with sulfide has not been previously demonstrated in the PNSB *Phaeospirillum* (47, 60).

Instead, we propose that sulfide oxidation may serve as a detoxification mechanism. This may have enabled their relative success compared with other MAGs in the deeper sediments of the column, where sulfide is expected to be present at high concentrations (**Fig. 1**). Such a mechanism has been reported previously in the *Cyanobacteriota* (61), including, possibly, those we observed here. There is also evidence that some *Cyanobacteriota* can use sulfide, when present at high concentrations, as an electron donor for anoxygenic photosynthesis (62–64). This could explain the dominance of *Cyanobacteriales* Bin.017_ZZ in the sediments of the Winogradsky column, where oxygen may be more limited while sulfide is abundant (**Fig. 1**).

Some of the MAGs may further oxidize ZVS to sulfite (**Fig. 2**). The two GSB MAGs, *Phaeospirillum* Bin.004_BAB, and *Chromatiaceae* Bin.008_AM may use the reverse dissimilatory sulfite reductase (rDsrAB), while *Kamptonema* Bin.017_EG might use sulfur dioxygenase (Sdo). For most of these sulfur oxidizers, sulfite appears to be the endpoint, though *Chlorobaculum* Bin.017_DR encodes a thiosulfate reductase (Phs) homolog that might operate in reverse to produce thiosulfate from sulfite (60). The deep-sediment-dominant *Rhodopseudomonas* Bin.015_MK may have the greatest flexibility in oxidizing sulfur compounds, encoding both FccB and a complete Sox system, which may allow it to use sulfide, sulfite, and thiosulfate as electron donors (**Fig 2**). While described as PNSB, *Rhodopseudomonas* have well-characterized sulfur-oxidation capabilities (61–64).

Of note, we failed to recover any MAGs of sulfate-respiring bacteria (SRB). However, SRB are necessary to transform the sulfate added to the column into reduced sulfur. Instead, we performed a read-based taxonomic analysis, which revealed several lower-abundance SRB families—particularly *Desulfovibrionaceae, Desulfobulbaceae,* and *Desulfobacteraceae*—within the column sediments (**Table S8**) that may facilitate this reduction step.

### Complex carbon degradation pathways are widely distributed among MAGs

Winogradsky columns offer an opportunity to analyze complex carbon metabolism across redox-stratified microbiota. Here, the shredded coffee filter added to the column is expected to be composed mostly of cellulose, with some hemicellulose (xylans and mannans) (70). Additionally, pond water added to the top of the column contained leaf and stick litter, contributing lignin and pectin (71). The depolymerization of these polymers is carried out by functionally diverse carbohydrate-active enzymes (CAZymes) (67), thereby making these recalcitrant forms of carbon more accessible to the entire column community (68). We found CAZymes distributed across metabolically and taxonomically diverse MAGs, allowing for many possible pathway configurations for the complete depolymerization of these substrates (**Fig. 3, Fig. S3, Table S9**).

**Figure 3.**
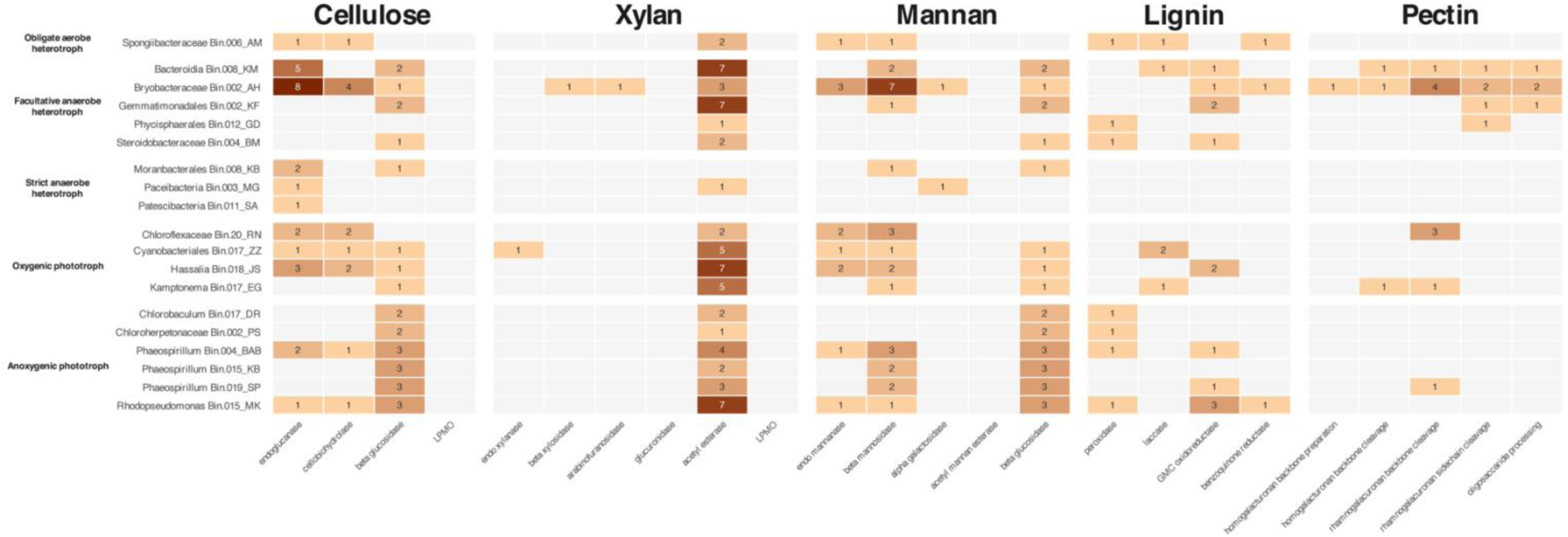
Distributions of CAZymes associated with lignocellulosic degradation across the MAGs. Counts show the number of hits per MAG. The CAZy families associated with each enzyme are listed in **Table S10. Figure S2** shows the distribution of these individual families across the MAGs.

However, a recent study examining CAZymes in metagenomic and metaproteomic datasets from freshwater sediment enrichments suggested that most depolymerization is carried out by a small number of specialists adapted to the specific lignocellulose source (72). Thus, this genomic capacity may not always be fully active.

Hits for CAZyme families associated with the depolymerization of cellulose, mannan, and lignin are widely distributed across all MAGs (**Fig. 3**). However, CAZyme families associated with xylan depolymerization (except for acetyl esterases) are more uncommon, with one endo-xylanase (GH10) encoded by the oxygenic phototroph *Cyanobacteriales* Bin.017_ZZ and one beta-xylosidase/arabinofuranosidase (GH43) encoded by the facultatively anaerobic heterotroph *Bryobacteraceae* Bin.002_AH (**Fig. 3**). This lack of xylan-related genes is notable as the coffee filter is softwood-derived paper, which typically contains 5-7% xylans (70). Thus, it is interesting that the only endo-xylanase is encoded by the very dominant *Cyanobacteriales* Bin.017_ZZ (**Figs. 1, S2**), possibly privileging this population with access to an otherwise poorly exploited organic carbon pool. In contrast, acetyl esterases (CE1-6), which deacetylate xylan, are ubiquitous across the genomes (**Fig. 3**). Some of these acetyl esterases are also known to deacetylate other substrates like chitin, peptidoglycan, and lignin (72, 73). Such deacetylation liberates acetate, providing an additional pool of labile carbon to other community members (74). CAZymes associated with pectin depolymerization are similarly restricted to a smaller subset of the community. The facultatively anaerobic heterotrophs *Bryobacteraceae* Bin.002_AH and *Bacteroidia* Bin.008_KM are the only MAGs encoding a complete or nearly complete pectin depolymerizing suite. They are also the only MAGs encoding CAZymes (PL1 and PL11) associated with the cleavage of the homogalacturonan and rhamnogalacuronan backbone of pectin. In general, these two MAGs have the most comprehensive suite of lignocellulosic CAZymes.

Depolymerization products, such as oligosaccharides and organic acids, that are not assimilated into the biomass of primary degraders are predicted to be fermented by organisms with more restricted catabolic capacities. Among these secondary consumers are the *Patescibacteria*, which were substantive members of the deeper subsurface community. *Paceibacteria* Bin.003_MG and *Moranbacterales* Bin.008_KB accounted for approximately 6% and 2% of the community in this region, respectively (**Fig. 1, Fig. S2**), while a third MAG (*Patescibacteria* Bin.011_SA) occurred at lower abundance in the biofilms of column 2 as well as the shallow subsurface sediments. The ultrasmall *Patescibacteria* are thought to play a key role in organic carbon cycling in natural environments by generating labile fermentation products that can be used by the broader community (76). Due to their reduced genomes, they also rely heavily on externally derived metabolites, particularly those from host cells (77). Consistent with this lifestyle, all three *Patescibacteria* MAGs encode a D-lactate dehydrogenase and lack respiratory machinery. Among the three, they encode multiple glycoside hydrolase (GH1, GH74, GH36) CAZymes that may facilitate the utilization of shorter-chain products generated during lignocellulose depolymerization (**Fig. 3**), potentially funneling this organic carbon towards lactate production. These MAGs lack complete versions of the upper glycolytic pathway, the pentose phosphate pathway, and the TCA cycle—like other *Patescibacteria* (78). Some *Patescibacteria* might utilize a truncated glycolytic pathway, involving a partial pentose phosphate pathway, to support fermentation (79). This route may be possible in *Patescibacteria* Bin.003_MG and *Patescibacteria* Bin.011_SA; however, *Moranbacterales* Bin.008_KB lacks any pentose phosphate pathway genes.

Host-associated biomass turnover may provide a second route by which *Patescibacteria* acquire labile carbon. Like other *Patescibacteria*, these MAGs are expected to be ectosymbiotic (76), a prediction supported in part by the presence of genes encoding a Type IV pilus (**Table S11**), which has been shown to be involved in host cell attachment (77). All three MAGs also encode M23B-family endopeptidases, a family that includes enzymes capable of hydrolyzing bacterial peptidoglycan (78, 79). *Moranbacterales* Bin.008_KB similarly encodes endopeptidase NlpC/P60, which can also lyse bacterial cell walls (80). Additional peptidases identified in the genome annotations may further facilitate the acquisition of amino acids from host-derived proteins. *Moranbacterales* Bin.008_KB and *Patescibacteria* Bin.011_SA also encode malic enzyme, providing a route for host-derived malate to enter metabolism and support lactate fermentation, as recently proposed for other *Patescibacteria* (81). Thus, we suggest a model in which *Patescibacteria* inhabiting the Winogradsky column transform plant-derived organic carbon, both directly and indirectly (via scavenging of host biomass ultimately derived from the same carbon pool), into labile fermentation products that support the broader microbial community.

## CONCLUSIONS

The steep chemical gradients of the Winogradsky column sustain a microbial community dominated by largely uncultivated taxa, and these populations partition into different guilds that are predicted to divide the labor of carbon, sulfur, and nitrogen cycling. We suggest that Winogradsky columns make an interesting, tractable model for understanding microbial community metabolism in redox-stratified systems—one that also renders these otherwise invisible processes spatially perceptible. Finally, we hope these findings prove equally valuable to educators and their students, for whom reading about these processes can complement what the column lets them see unfold.

## AUTHOR CONTRIBUTIONS

**Conceptualization:** JLG, JY; **Data curation:** SPA, KPB, GTD, MJG, ZZ, KB, AJM, PS, JLG; **Formal analysis:** SPA, KPB, GTD, MJG, ZZ, JLG; **Investigation:** SPA, KPB, GTD, MJG, ZZ, KB, AJM, PS, BB, RF-N, KF, EG, AGH, MGK, BEM, KM, SP, DR, JLS, CARB; **Methodology:** SPA, KPB, GTD, MJG, ZZ, CARB, JY, JLG; **Resources:** JLG, JY; **Supervision:** JLG; **Validation:** JLG; **Visualization:** KPB, GTD, JLG**; Writing – original draft:** SPA, KPB, GTD, MJG, ZZ, CARB, JLG; **Writing – review & editing:** SPA, KPB, GTD, MJG, ZZ, KB, AJM, PS, BB, RF-N, KF, EG, AGH, MGK, BEM, KM, SP, DR, JLS, CARB, JY, JLG

## ACKNOWLEDGEMENTS

This work was funded by MICROnet ⏤ Microbiomes In Computational Research Opportunities Network (NSF # 2418285). We are also grateful for the Landscape Architecture graduate students at SUNY ESF who established the Winogradsky columns as part of their coursework.

