## Supporting Methods, Tables S10-S11, Figs. S1-S3 for "Genome-resolved insights into microbial diversity and elemental cycling in Winogradsky columns"

**SUPPORTING INFORMATION**

**ADDITIONAL METHODOLOGICAL DETAILS**

***DNA extraction and sequencing.*** To sample the sediments and biofilms from the Winogradsky columns, first, water was drained from the columns. About 250 mg of sediment was “cored” from the edges (*i.e.*, the light-exposed regions) using a sterilized spatula. We note that sampling from the lower sites was particularly difficult, and little pieces of biofilm from the higher sections of the column were sometimes collected as the spatula was withdrawn. DNA extractions were performed using the ZymoBIOMICS DNA Miniprep kit, following the protocol with the following changes: 1) Lysis was performed using the Vortex Genie for 40 minutes of continuous bead beating. 2) Eluted DNA was not run through the Zymo-Spin III-HRC Filter. Five samples with DNA concentrations over 10 ng/μL were sent to SeqCoast Genomics (Portsmouth, NH) for sequencing. Library preparation was performed using the Illumina DNA Prep tagmentation kit with Illumina Unique Dual Indexes. Sequencing was performed on the Illumina NextSeq2000 platform to generate 2 x 150 bp paired end reads. Demultiplexing, quality filtering, and read trimming were performed using DRAGEN (v4.2.7) (1).

***Metagenomic assembly, binning, and annotation:*** Metagenomic assembly and binning of metagenome-assembled genomes (MAGs) were performed using the US Department of Energy (DOE) KnowledgeBase (KBase) platform (2) using default settings. Read quality was verified using FastQC (v0.12.1). The five metagenomic libraries were assembled and binned both separately and as a single co-assembly to maximize MAG recovery. Metagenomes were assembled using metaSPAdes (v3.15.3) (3). Binning was performed using CONCOCT (v1.1) (4), MaxBin2 (v2.2.4) (5), MetaBAT2 (v1.7) (6), and COCACOLA (v1.0) (7). The bin outputs were then optimized using DAS Tool (v1.1.2) (8). Taxonomic assignment was performed using GTDB-tk with GTDB r214 (9). For the three *Phaeospirillum* MAGs, average nucleotide identity was computed using FastANI (v0.1.3)(10). Default settings were used for all the above analyses.

Genome completeness and contamination were checked using CheckM (v1.10.18) (11) in KBase. For members of the *Patescibacteria*, we estimated these metrics using phylum-specific markers previously described (12). Barrnap (v1.10.5) (<https://github.com/tseemann/barrnap>) was used to predict 16S and 5S rRNA gene sequences. tRNAscan-SE (v2.0.12) was used to predict tRNA gene sequences (13). We used the MIMAG standards for reporting MAG quality (14), retaining all medium- and high-quality draft genomes for further analysis. High-quality draft genomes were defined as: >90% completion, <5% contamination, and the presence of the 23S, 16S, and 5S rRNA genes and at least 18 tRNA genes. Medium-quality draft genomes were defined as: ≥50% completion and <10% contamination. For the purposes of our analysis, we also retained two MAGs with >40% (but <50%) completion and <5% contamination, and one MAG with >90% completion and <20% (but>10%) contamination. In addition, we used GUNC (v1.1.1), with default parameters, as a complementary assessment of genomic chimerism (15). With the retained MAGs, CoverM (v0.7.0) was used *(--mapper strobealign, --min-read-percent-identity 95, --min-read-aligned-percent 75*) to calculate transcripts per million (TPM) and relative abundance of the MAGs across the five sequencing libraries (16). For downstream analyses, the abundance data from the two replicate samples from the surface biofilm in column 1 were averaged.

Annotations were performed on KBase. MAGs were annotated using RASTtk (v1.073) (17) and DRAM (v0.1.2) (18). Further annotation was performed to identify MAGs encoding carbohydrate-active enzymes (CAZymes) involved in lignocellulose depolymerization. CAZy family hits identified by DRAM were associated with each enzymatic degradation step of the main lignocellulosic polymers (cellulose, xylan, mannan, lignin, and pectin), as defined by the CAZy database (**Table S10**) (19). Further metabolic profiling was performed using a set of Hidden Markov Models (HMMs) from the environmental bioelement MicroTrait collection (20) with searches implemented using HMMER (v3.3.2) (21). Default parameters were used for all analyses. Type IV pilus proteins were identified in the *Patescibacteria* MAGs by performing a BLASTp search (22) against homologs from *Candidatus Chazhemtobacterium aquaticus.* Positive determination included greater than 25% identity and greater than 70% alignment against the query protein.

**SUPPLEMENTAL TABLES AND FIGURES**

| **Table S10.** Enzymatic activities and substrates associated with the CAZy families analyzed here | | |
| --- | --- | --- |
| **Polymer** | **Enzymatic Degradation Step** | **CAZy Families** |
| Cellulose | endoglucanase | GH5, GH9, GH12, GH44, GH45, GH48, GH74 |
|  | cellobiohydrolase | GH5, GH9, GH48 |
|  | beta glucosidase | GH1, GH3 |
|  | lytic polysaccharide monooxygenase | AA10 |
| Xylan | endo xylanase | GH10, GH11, GH30 |
|  | beta xylosidase | GH43, GH52, GH54 |
|  | arabinofuranosidas | GH43, GH51, GH54, GH62 |
|  | glucuronidase | GH67, GH115 |
|  | acetyl esterase | CE1, CE2, CE3, CE4, CE5, CE6 |
|  | lytic polysaccharide monooxygenase | AA14 |
| Mannan | endo mannanase | GH5, GH26, GH113 |
|  | beta mannosidase | GH1, GH2, GH5 |
|  | alpha galactosidase | GH27, GH36 |
|  | acetyl mannan esterase | CE2 |
|  | beta glucosidase | GH1, GH3 |
| Lignin | peroxidase | AA2 |
|  | laccase | AA1 |
|  | GMC oxidoreductase | AA3 |
|  | benzoquinone reductase | AA6 |
| Pectin | homogalacturonan backbone preparation | CE8, CE12, CE13 |
|  | homogalacturonan backbone cleavage | GH28, PL1, PL2, PL3, PL9, PL10 |
|  | rhamnogalacuronan backbone cleavage | GH28, PL4, PL11, PL26, GH78, GH106 |
|  | rhamnogalacuronan sidechain cleavage | GH34, GH51, GH54, GH62, GH93, GH53, GH30, GH35, GH105, GH88 |
|  | oligosaccaride processing | GH105, GH88 |

| Table S11. Surface-adhering type IV pilus proteins identified in the *Patescibacteria* MAGs | | | | | |
| --- | --- | --- | --- | --- | --- |
|  | Multimodular transpeptidase-transglycosylase | Twitching motility protein PilT | Type IV fimbrial assembly protein PilC | Type IV fimbrial assembly, ATPase PilB | Type IV pilus Assembly Protein PilA |
| *Patescibacteria* Bin.003_MG | ✔ | ✔ | ✔ | ✔ | ✔ |
| *Moranbacterales* Bin.008_KB | ✔ | ✔ | ✔ | ✔ | ✔ |
| *Patescibacteria* Bin.011_SA | ✔ | X | X | ✔ | X |

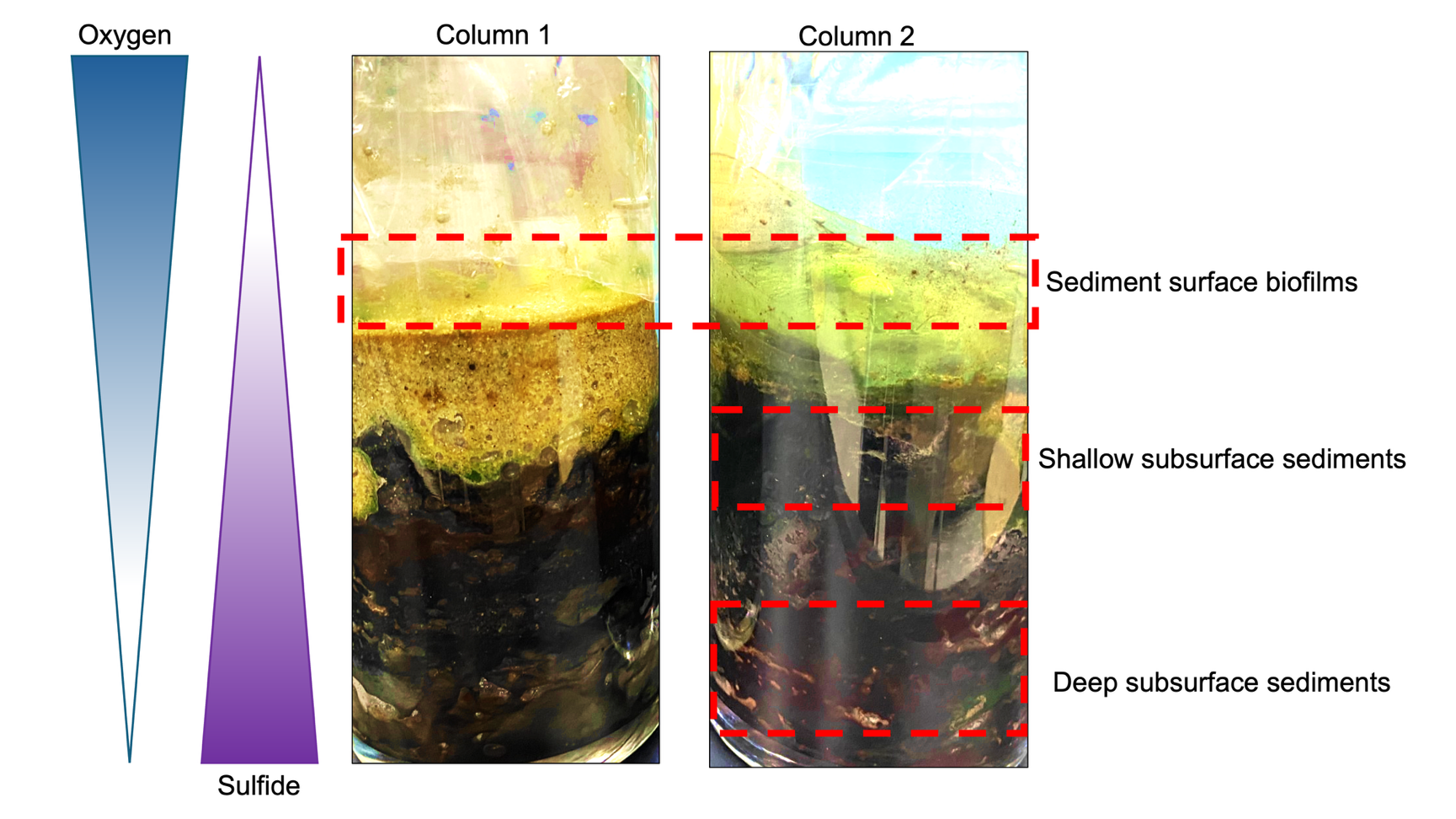
**Fig. S1: Images of the two Winogradsky columns.** Oxygen and sulfide gradients, inferred from (23, 24), are depicted to the left. Regions where DNA was extracted are indicated by the red dashed-line boxes. Note that the brightness and the saturation of the lower-quality cellphone camera images were enhanced to make the darker red and green pigmentation more visible for the purposes of appreciating the layers referenced in the paper.

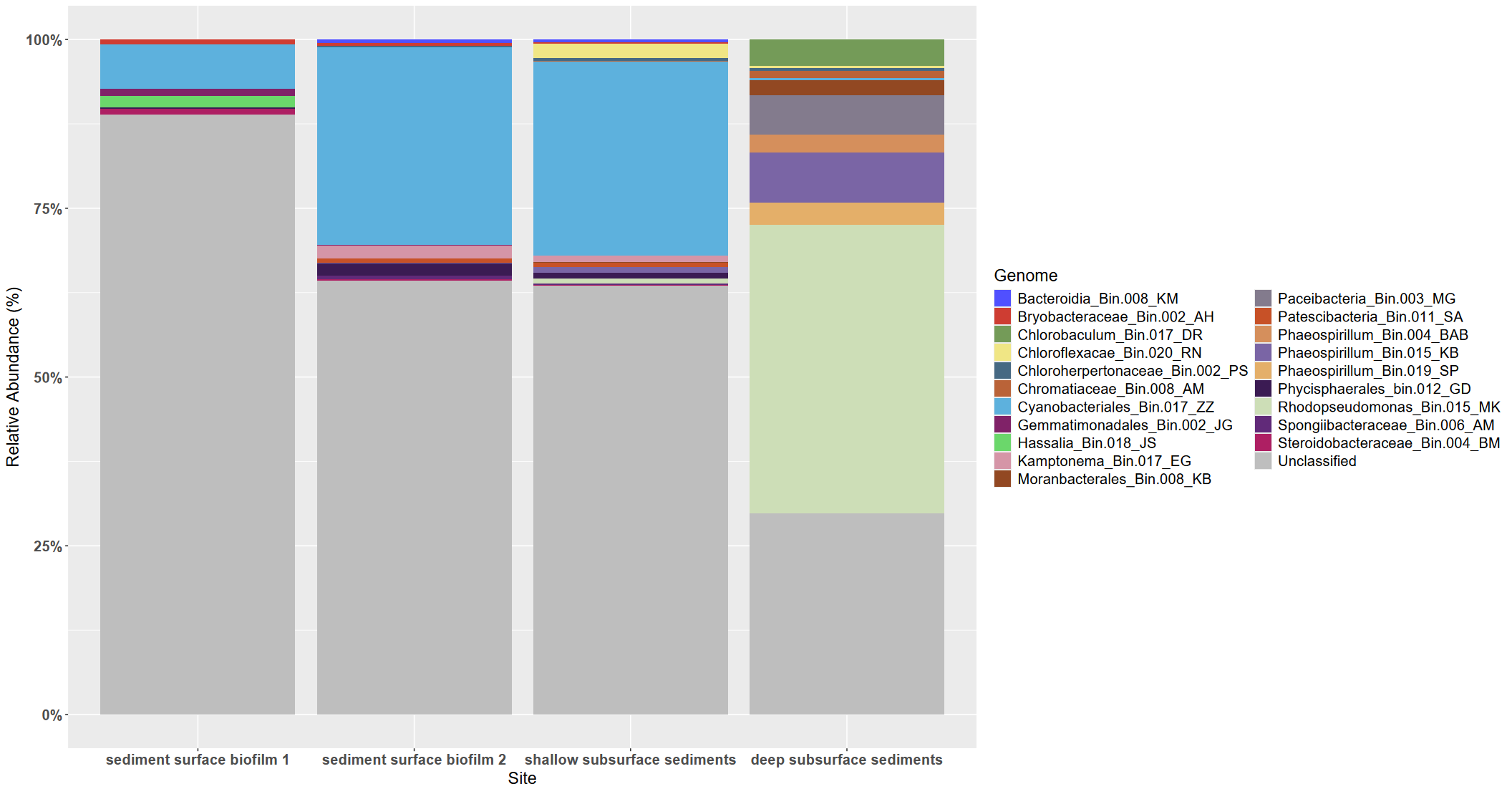
**Fig. S2. Relative abundance of MAGs** **within the Winogradsky column community.** Abundances are estimated from the proportion of reads in each sample that mapped to a particular MAG. Images of the columns are shown in **Fig. S1**. “Unclassified” reads did not map to any of the MAGs.

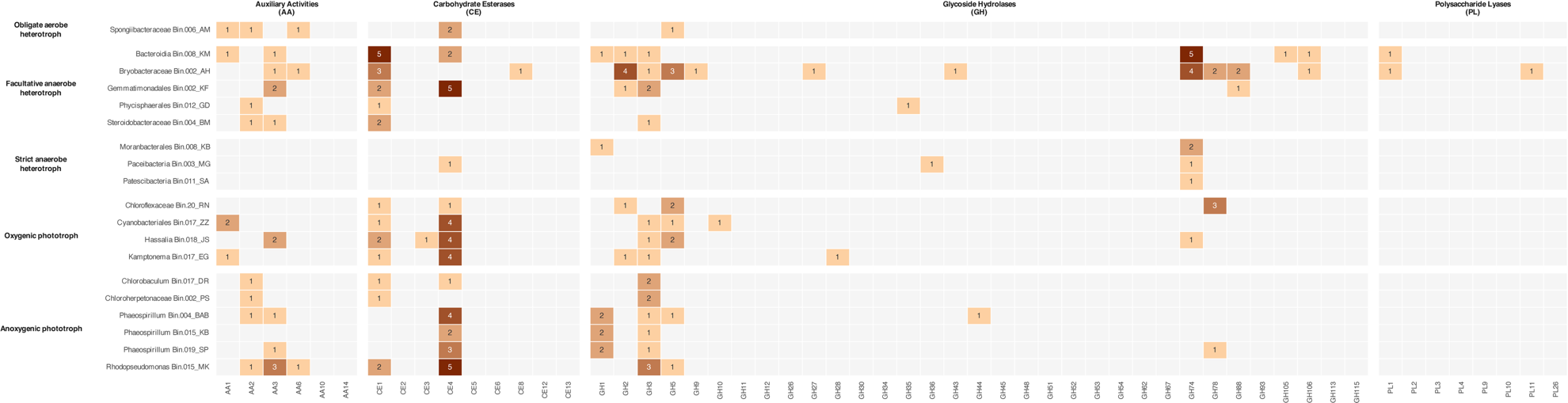

**Figure S3. Individual CAZy family hit associated with lignocellulosic degradation across the MAGs.** Counts show the number of hits per MAG. The enzymatic activities of the CAZy families are listed in **Table S10.**
